# A practical guide for inferring reliable dominance hierarchies and estimating their uncertainty

**DOI:** 10.1101/111146

**Authors:** Alfredo Sánchez-Tójar, Julia Schroeder, Damien R. Farine

**Author notes:** corresponding authors; complete postal address: Max Planck Institute for Ornithology, Eberhard-Gwinner-Straße, 82319, Seewiesen, Germany.; complete postal address: Fachbereich Biologie, Universität Konstanz, Universitätsstraße 10, 78464, Konstanz, Germany.

## Abstract

Many animal social structures are organized hierarchically, with dominant individuals monopolizing resources. Dominance hierarchies have received great attention from behavioural and evolutionary ecologists. As a result, there are many methods for inferring hierarchies from social interactions. Yet, there are no clear guidelines about how many observed dominance interactions (i.e. sampling effort) are necessary for inferring reliable dominance hierarchies, nor are there any established tools for quantifying their uncertainty. In this study, we simulated interactions (winners and losers) in scenarios of varying steepness (the probability that a dominant defeats a subordinate based on their difference in rank). Using these data, we (1) quantify how the number of interactions recorded and hierarchy steepness affect the performance of three methods, (2) propose an amendment that improves the performance of a popular method, and (3) suggest two easy procedures to measure uncertainty in the inferred hierarchy. First, we found that the ratio of interactions to individuals required to infer reliable hierarchies is surprisingly low, but depends on the hierarchy steepness and method used. We then show that David’s score and our novel randomized Elo-rating are the two best methods, whereas the original Elo-rating and the recently described ADAGIO perform less well. Finally, we propose two simple methods to estimate uncertainty at the individual and group level. These uncertainty measures further allow to differentiate non-existent, very flat and highly uncertain hierarchies from intermediate, steep and certain hierarchies. Overall, we find that the methods for inferring dominance hierarchies are relatively robust, even when the ratio of observed interactions to individuals is as low as 10 to 20. However, we suggest that implementing simple procedures for estimating uncertainty will benefit researchers, and quantifying the shape of the dominance hierarchies will provide new insights into the study organisms.

**Highlights:**

- David’s score and the randomized Elo-rating perform best.
- Method performance depends on hierarchy steepness and sampling effort.
- Generally, inferring dominance hierarchies requires relatively few observations.
- The R package “aniDom” allows easy estimation of hierarchy uncertainty.
- Hierarchy uncertainty provides insights into the shape of the dominance hierarchy.

## INTRODUCTION

Many animal social structures are organized hierarchically, with some individuals – the dominants – monopolizing resources and therefore presumably monopolizing fitness, too. First described in the domestic fowl (Schjelderupp-Ebbe, 1922), dominance hierarchies have received great attention from empiricists and theoreticians in behavioural and evolutionary ecology. Dominance hierarchies have widely been described in insects (Choe, 1994), fishes (Polačik & Reichard, 2009), reptiles (Bush, Quinn, Balreira, & Johnson, 2016), birds (Devost, Jones, Cauchoix, Montreuil-Spencer, & Morand-Ferron, 2016) and mammals (Majolo, Aureli, & Schino, 2012), including humans (von Rueden, Gurven, & Kaplan, 2008). Extensive theoretical efforts have been made on understanding how dominance hierarchies are formed and maintained (e.g. Dugatkin & Earley, 2004; Parker, 1974; Sasaki et al., 2016).

The importance and prevalence of dominance hierarchies in nature (reviewed in Drews, 1993) has led to the development of many methods for inferring dominance hierarchies from social interactions (reviewed in Bayly, Evans, & Taylor, 2006; Briffa et al., 2013; de Vries, 1998; Whitehead, 2008), yet, clear guidelines for inferring reliable dominance hierarchies are still missing. These methods can be classified into those estimating the rank of the individuals (i.e. an ordinal score: 1,2,…,n; e.g. I&SI: de Vries, 1998; ADAGIO: Douglas, Ngonga Ngomo, & Hohmann, 2017) and those estimating non-integer indices of success from which individuals can further be ranked if required (e.g. David's score: David, 1987; Elo-rating: Elo, 1978; Fig. 1). Contrary to ranks, indices have the advantage of allowing parametric statistical testing. Index-generating methods are further classified into those based on interaction matrices (e.g. David’s score) and those based on the temporal sequence of interactions (e.g. Elo-rating; Fig. 1). What all methods have in common is that they require researchers to record agonistic dyadic interactions among individuals as input data. Despite great research effort, there are surprisingly few clear guidelines about how many interactions are needed for inferring reliable dominance hierarchies, nor are there any established tools for quantifying the uncertainty of a dominance hierarchy (i.e. determine whether sufficient observations were made).

**Figure 1.**
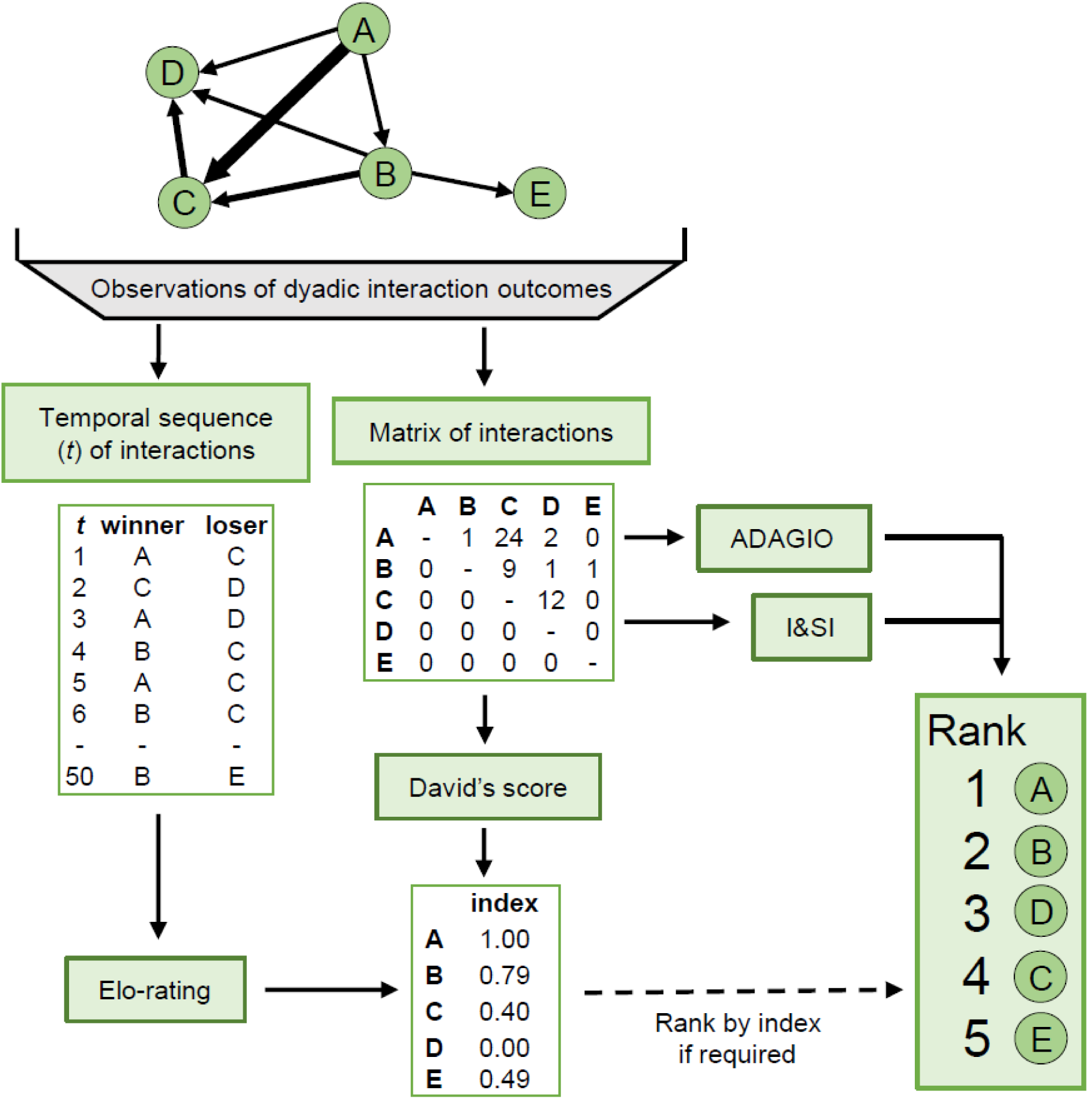
Diagram highlighting the different steps required to infer dominance hierarchies. First, the outcomes of dyadic agonistic interactions between individuals are recorded either in the form of a matrix or as a temporal sequence of winners and losers. Second, different methods can be used to infer either individual ranks or individual non-integer indices of success, which can further be used to rank the individuals and therefore to infer the dominance hierarchy of the group.

One source of uncertainty is the steepness of the dominance hierarchy. Dominance hierarchies can range from very steep, where dominant individuals win all conflicts, to completely flat, non-existent hierarchies, where dyadic outcomes are unpredictable. These different scenarios, which are *a priori* unknown by the researcher, are expected to affect the performance of the method inferring hierarchies. Several efforts have been made to assist researchers selecting an appropriate method, however, most of these attempts focused on comparing the level of agreement among several methods when applied to real datasets (e.g. Balasubramaniam et al., 2013; de Vries, 1998; Gammell, de Vries, Jennings, Carlin, & Hayden, 2003; Neumann et al., 2011). The problem with using real datasets is that the real steepness of the hierarchy and the real rank of the individuals are unknown. Thus, while cases where two or more methods closely match one-another could signify that they are robust, this could also mean that they suffer from a common bias, and such comparisons provide no information about their accuracy. A better approach is to (i) simulate artificial datasets containing individuals of known rank, (ii) simulate interactions among those individuals under different scenarios of known steepness, and (iii) then test the validity of the method(s) by correlating the inferred hierarchy to the original known hierarchy from step (i). This approach further allows estimating the degree of uncertainty of the inferred ranks, and how this varies based on the steepness of the hierarchy and the number of interactions observed.

An additional source of uncertainty is the skewness in the propensity for individuals to interact. As with many other biological processes that generate count data, the number of interactions per individual often follow a Poisson distribution. This means that few individuals have many interactions, whereas most individuals have few interactions. The unequal distribution of interactions leads to interaction datasets that are sparse. Sparse datasets are very common in the dominance literature (McDonald & Shizuka, 2013) and sparseness can potentially affect the performance of the method (de Vries, 1998; Gammell et al., 2003; Neumann et al., 2011). Further, such distribution could over-inflate the perceived quality of an interaction dataset that in fact contains too few interactions to estimate most individuals’ ranks.

There is increasing awareness of the need to estimate uncertainty of social data (e.g. Farine & Strandburg-Peshkin, 2015; Lusseau, Whitehead, & Gero, 2008). More than a decade ago Adams (2005) proposed a Bayesian approach to estimate hierarchy uncertainty, however, behavioural and evolutionary ecologists have not yet broadly adopted Bayesian procedures, possibly due to their apparent complexity. Thus, uncertainty is still rarely measured when studying dominance hierarchies (but see Kelstrup, Hartfelder, & Wossler, 2015; Sheppard et al., 2013). Further, estimating uncertainty is often seen as a step needed simply to ‘tick a box’. Here we demonstrate that estimating uncertainty can be simple to implement, and that doing so can also generate new insights into the processes being studied.

In this study, we simulated artificial datasets and interactions under a wide range of scenarios, from very steep to completely flat, non-existent hierarchies, to: (1) quantify how the number of interactions recorded (sampling effort) and hierarchy steepness affect the performance of different methods inferring the correct hierarchy, (2) propose an amendment to a popular method, the original Elo-rating, to improve its performance, and (3) suggest two easy procedures to measure the uncertainty of the inferred hierarchies. We focus our study on three index-generating methods (plus our method) that have been commonly used in the recent literature. First, we evaluate David’s score (David, 1987), a widely used matrix-like method that is based on the paired comparisons paradigm (e.g. Jennings, Carlin, & Gammell, 2009; Rat, van Dijk, Covas, & Doutrelant, 2015). Second, we evaluate the original Elo-rating (Elo, 1978), a sequential method that is becoming popular in the study of animal behaviour (e.g. Franz, Mclean, Tung, Altmann, & Alberts, 2015; Snyder-mackler et al., 2016; Strandburg-Peshkin, Farine, Couzin, & Crofoot, 2015). Following our evaluation of this method, we suggest a modification to the original Elo-rating that improves estimates and provides a measure of uncertainty for each individual’s rank. Finally, we evaluate ADAGIO, a recently described method that is based on the extraction and graphical representation of directed acyclic graphs (Douglas et al., 2017). From our results, we derive recommendations on the sampling effort required to infer reliable dominance hierarchies.

## MATERIALS AND METHODS

Our general approach consisted of (i) generating artificial datasets containing individuals of known rank, (ii) simulating interactions among those individuals under different hierarchy scenarios of known steepness and propensities to interact, and (iii) testing the performance of the different methods under the different scenarios.

We implemented all of our simulations in R v.3.3.2 (R Core Team, 2016; see Sánchez-Tójar, Schroeder, & Farine, 2017). We created a user friendly R package named “aniDom” to infer dominance hierarchies using the original and the randomized Elo-rating method. The package also allows estimating hierarchy uncertainty (see below) as well as plotting the steepness of the hierarchy. We used the R package “EloRating” to calculate David’s scores (Neumann & Kulik, 2014) and a separate software for the ADAGIO method (Douglas et al., 2017).

### i Generating artificial groups of individuals

We generated an artificial dataset containing 50 individuals whose ranks were sequentially assigned from 1 to 50 (i.e. linear hierarchy). Because count data (such as the number of interactions) typically follow a Poisson distribution (Zuur, Ieno, Walker, Saveliev, & Smith, 2009), we used a Poisson process to generate a varying propensity for each individual to interact. Other distributions gave qualitatively similar conclusions (Supplementary Material 1). Throughout we defined the “ratio of interactions to individuals” as the number of interactions (d, where one interaction is between two individuals) divided by the total number of individuals (N), that is 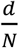, rather than the arithmetic mean of the number of interactions per individual (*i*), that is *d̄_l_,* which could be twice the value we report. For example, in a dataset containing 10 interactions between two individuals, 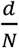 would equal 5, whereas *d̄_l_* would equal 10. We repeated all analyses with datasets containing 10 individuals (Supplementary Material 2).

### ii. Simulating interactions within the group

We generated simulated datasets. In each interaction dataset, the outcome of the dyadic interactions was determined by the specific hierarchy scenario implemented. We used a probabilistic approach to generate a wide range of hierarchy scenarios of different steepness. Specifically, we modelled the expected probability of winning for the higher ranked individual (P) following the equation:

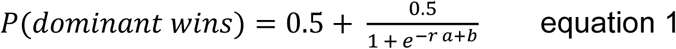

where *r* is the absolute difference in rank between the two individuals divided by the maximum absolute difference in rank possible in the dataset (i.e. 50 – 1 = 49 in our datasets), and thus, 0 < *r* ≤ 1. a and *b* are the values that determine the steepness of the hierarchy. For example, if *b* = −5 and a ≥ 0, the expected probability of winning for the higher ranked individual is essentially 1 at all times, regardless of the difference in rank between the two individuals, and thus this hierarchy is very steep (Fig. 2a). By contrast, if *b =* 35 and 0 ≤ *a* ≤ 30, the expected probability of winning for an individual is essentially always 0.5, regardless of the difference in rank between the two contestants, and thus there is no hierarchy (Fig. 2i). Figure 2 shows a wide range of the parameter space of equation 1, i.e. it shows different possible hierarchy scenarios.

**Figure 2.**
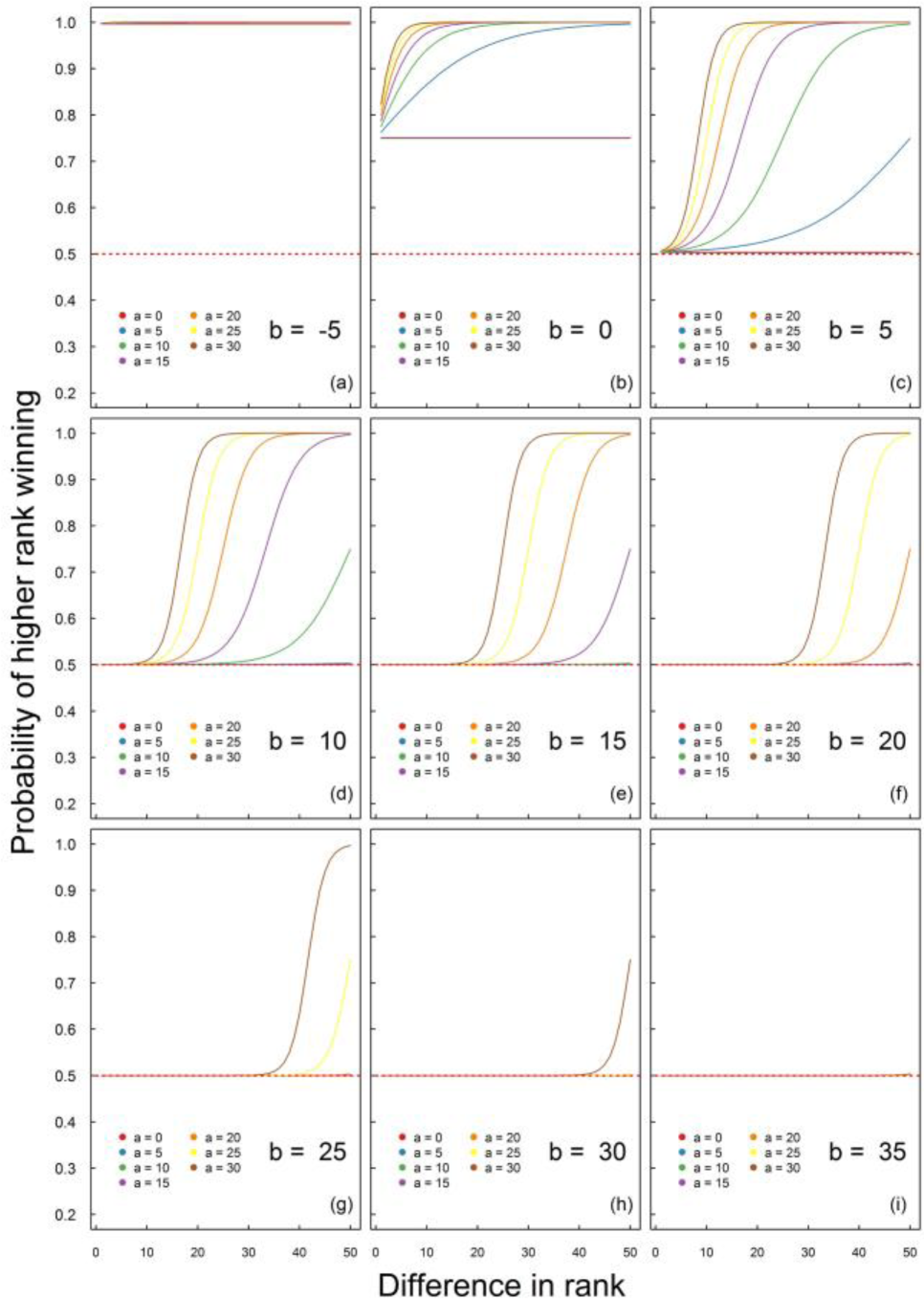
Parameter space of equation 1 showing a wide range of possible hierarchy scenarios that depend on the values assigned to *a* and *b*. Overall, panels are sorted from steep (panel a) to non-existent hierarchies (panel i). For each panel, steepness increases with *a*. The red dashed line shows where the probability of winning for the higher ranked individual (i.e. P(dominant wins)) equals 0.5, which corresponds to scenarios where dominance rank does not affect the probability of winning an interaction.

### iii. Tests

#### iii.a. Performance under different scenarios

To quantify the combined effects of sampling effort and hierarchy steepness on the ability to infer reliable dominance hierarchies, we inferred dominance hierarchies from the simulated data using the original Elo-rating, and compared these to the known simulated hierarchies. We explored a total of 42 hierarchy scenarios of different steepness. Specifically, we examined the following *b* values of equation 1: - 5, 0, 5, 10, 15 and 20. For each of those *b* values, we investigated the following *a* values: 0, 5, 10, 15, 20, 25 and 30. Hereafter, the initial Elo-rating for each individual was set to 1 000 and *k* (a parameter in the Elo-rating function) was set to 200 (for more details see Sánchez-Tójar et al. 2017). To quantify the relationship between sampling effort and the performance of the method, we assessed the performance for interactions datasets containing an increasing ratio of interactions to individuals of: 1, 4, 7, 10, 15, 20, 30, 40 and 50. For each scenario and ratio of interactions to individuals, we simulated 100 independent datasets, calculated individual ranks for each dataset following the original Elo-rating and obtained the Spearman rank correlation coefficient (hereafter rS) between the inferred and the known hierarchy. Therefore, if the hierarchies were identical, rS would equal 1.

#### iii.b. Comparing methods

We evaluated the performance of the four methods (David’s score, original Elo-rating, randomized Elo-rating and ADAGIO) under three hierarchy scenarios of intermediate steepness from equation 1, where *b* = 5, and *a* = 15, 10 and 5 (see Supplementary Material 3 for other scenarios). To quantify the relationship between sampling effort and the performance of the four methods, we assessed interactions datasets containing an increasing ratio of interactions to individuals of: 1, 4, 7, 10, 15, 20, 30, 40 and 50. For each scenario and ratio of interactions to individuals, we simulated 100 independent datasets, calculated individual ranks for each dataset following each of the four methods and, for each method, obtained the rS between the inferred and the known hierarchy.

#### iii.c. Randomized Elo-rating

One of the aims of the original Elo-rating is to enable tracking dynamic changes in rank over time. This means that the sequence in which interactions occur affects the inferred ranks. However, most behavioural studies assume that individual dominance rank is relatively stable over time (e.g. Poisbleau, Guillon, & Fritz, 2010). We propose an improvement of the original Elo-rating based on randomizing the order in which interactions occurred (*n* = 1 000 randomizations throughout). Randomising the order that interactions are recorded produces slightly different individual Elo-ratings each time, from which we can calculate a mean individual rank. This method also allows estimating the 95% range of individual ranks when run on a single interaction dataset. Note that in our method, we randomised all of the sequences of interactions because our data were simulated. However, in real datasets, one could test and account for underlying changes by randomising within certain periods of time, for example within each month, or test for winner-loser effects by randomising days but maintaining the order within each day and comparing these to full randomisations. Hereafter this new method is referred to as “randomized Elo-rating”, whereas its predecessor is referred to as “original Elo-rating”.

#### iii.d. Estimating hierarchy uncertainty

Using the randomizing Elo-rating, one can further estimate the repeatability of the *n* individual Elo-ratings. We explored repeatability using the same three scenarios described in section iii.c (see Supplementary Material 3 for other scenarios). Again, for each scenario and ratio of interactions to individuals, we simulated 100 independent interaction datasets, and calculated 1 000 individual Elo-ratings for each dataset following the randomized Elo-rating. We then calculated the repeatability of the individual Elo-ratings using the function rptGaussian() from the package ‘rptR’ 0.9.1.9000 (Schielzeth, Stoffel, & Nakagawa, 2016). We calculated repeatability based on Elo-ratings instead of ranks because, contrary to ranks, Elo-ratings approximately follow a Gaussian distribution.

Additionally, we tested another easy procedure to estimate hierarchy uncertainty which consists on splitting a dataset containing interactions into two halves, and then estimating the Spearman rank correlation coefficient between the two halves. We again investigated the same three hierarchy scenarios described in section iii.c (see Supplementary Material 3 for other scenarios). For each scenario and ratio of interactions to individuals, we simulated 100 independent interaction datasets, split each dataset in two, computed 1 000 individual ranks for each halve using the randomized Elo-rating, and calculated the r_S_ between the two inferred hierarchies.

## RESULTS

### Performance under different scenarios and sampling effort

We explored whether hierarchy steepness affects the performance of the method using the original Elo-rating. The performance of all four methods increased logarithmically with the ratio of interactions to individuals (Fig. 3 and 4). In most cases, the performance of the method reaches an asymptote after relatively few interactions. For very steep hierarchies, the original Elo-rating only needed a small ratio of interactions to individuals (10 or less) to satisfactorily infer the original hierarchy (e.g. Fig. 3a). In contrast, for non-existent hierarchies, the original Elo-rating visibly failed inferring the original hierarchy (e.g. *a ≤* 10 in Fig. 3c). Finally, for intermediate levels of steepness, the original Elo-rating had difficulties inferring the original hierarchy and needed a larger ratio of interactions to individuals to infer it reliably (e.g. *a* = 10 in Fig. 3b). However, in such intermediate scenarios, the improvements in the inferred hierarchy were only marginal beyond a ratio of interactions to individuals of 20.

**Figure 3.**
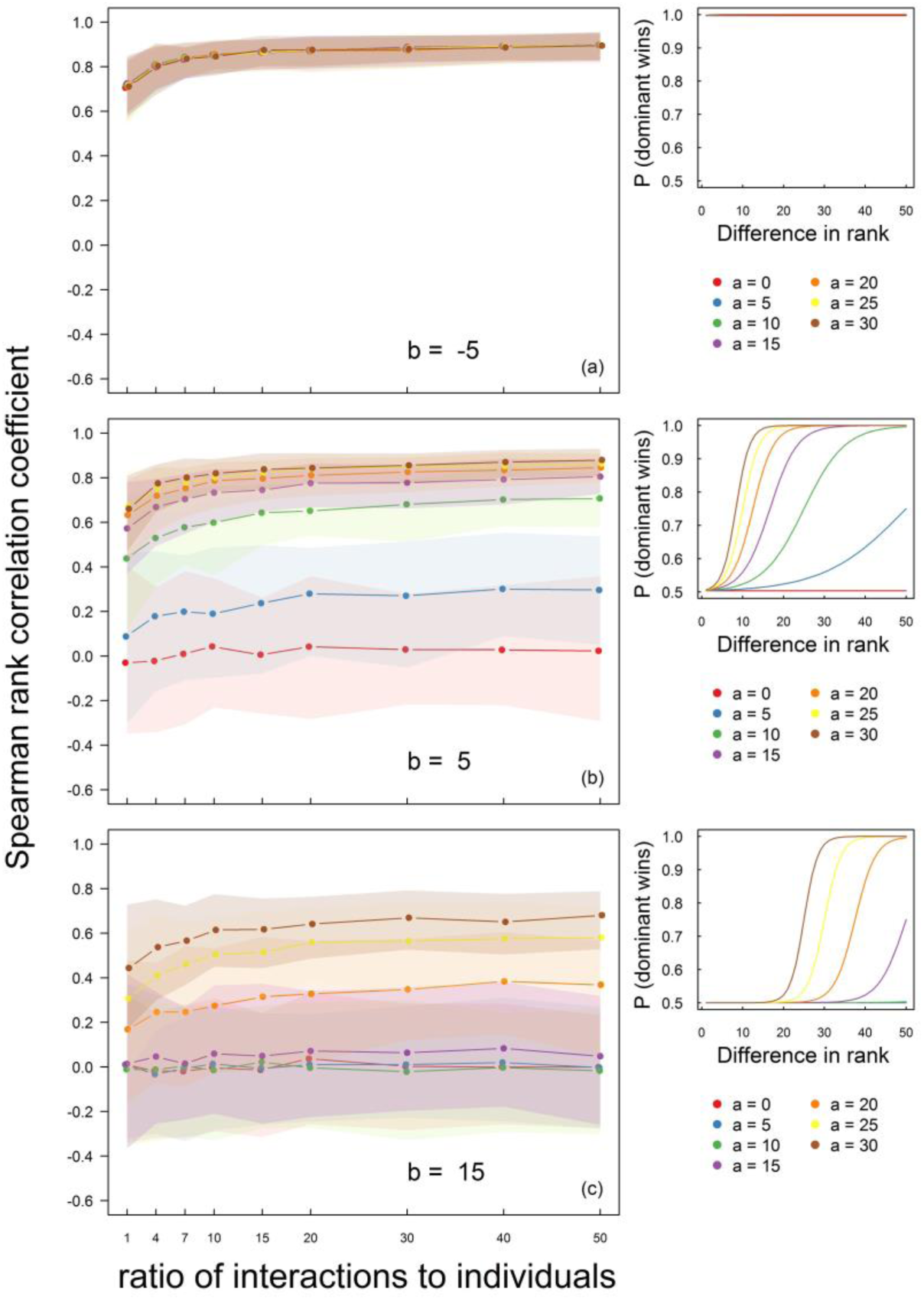
The performance of the original Elo-rating increases with the ratio of interactions to individuals and the steepness of the hierarchy. Solid lines and dots represent the mean Spearman rank correlation coefficient (r_S_) between the original and the inferred hierarchy; shading shows the 2.5% and 97.5% quartiles. The reduced right-hand side panels show the specific set of hierarchies simulated for generating the interaction datasets. Overall, panels are sorted from very steep (panel a) to flat hierarchies (panel c), and, for each panel, steepness increases with a. The different hierarchy scenarios shown were created following equation 1.

The steepness of the hierarchy not only affected the amount of data required to infer reliable dominance hierarchies, but also the overall ability to do so. In general, the method performed well even when closely-ranked individuals both often win contests. For example, even when the probability of the higher ranked individual winning was only ca 0.55 for a difference in rank of 10 (Fig. 3b, *a* = 15), the r_s_ between the original and the inferred hierarchy reached up to ca 0.70 when a ratio of interactions to individuals of 20 was recorded.

### Comparing methods

Next, we compared the performance of the four methods. For steep hierarchies, all four methods inferred similarly and satisfactorily the original hierarchy (Supplementary Material 3, Fig. S1a-c). Not surprisingly, for very flat hierarchies, the performance of all methods was extremely poor, even when a high ratio of interactions to individuals was recorded (Fig. 4c). At intermediate levels of steepness, we found that while all four methods had difficulties inferring the original hierarchy, the methods differed markedly in performance (Fig. 4a,b). Overall, David’s score performed best, closely followed by the randomized Elo-rating. By contrast, the original Elo-rating performed relatively poorly, which is probably because the sequence of the interactions, though it has little biological relevance, can have a large impact on the resulting ranks. The recently described ADAGIO did not perform well based on our simulated winner-loser data. Further, the shapes of the curves (when adding more interactions) across the methods were quite consistent, highlighting that it is unlikely that there are scenarios where their relative performances are flipped.

**Figure 4.**
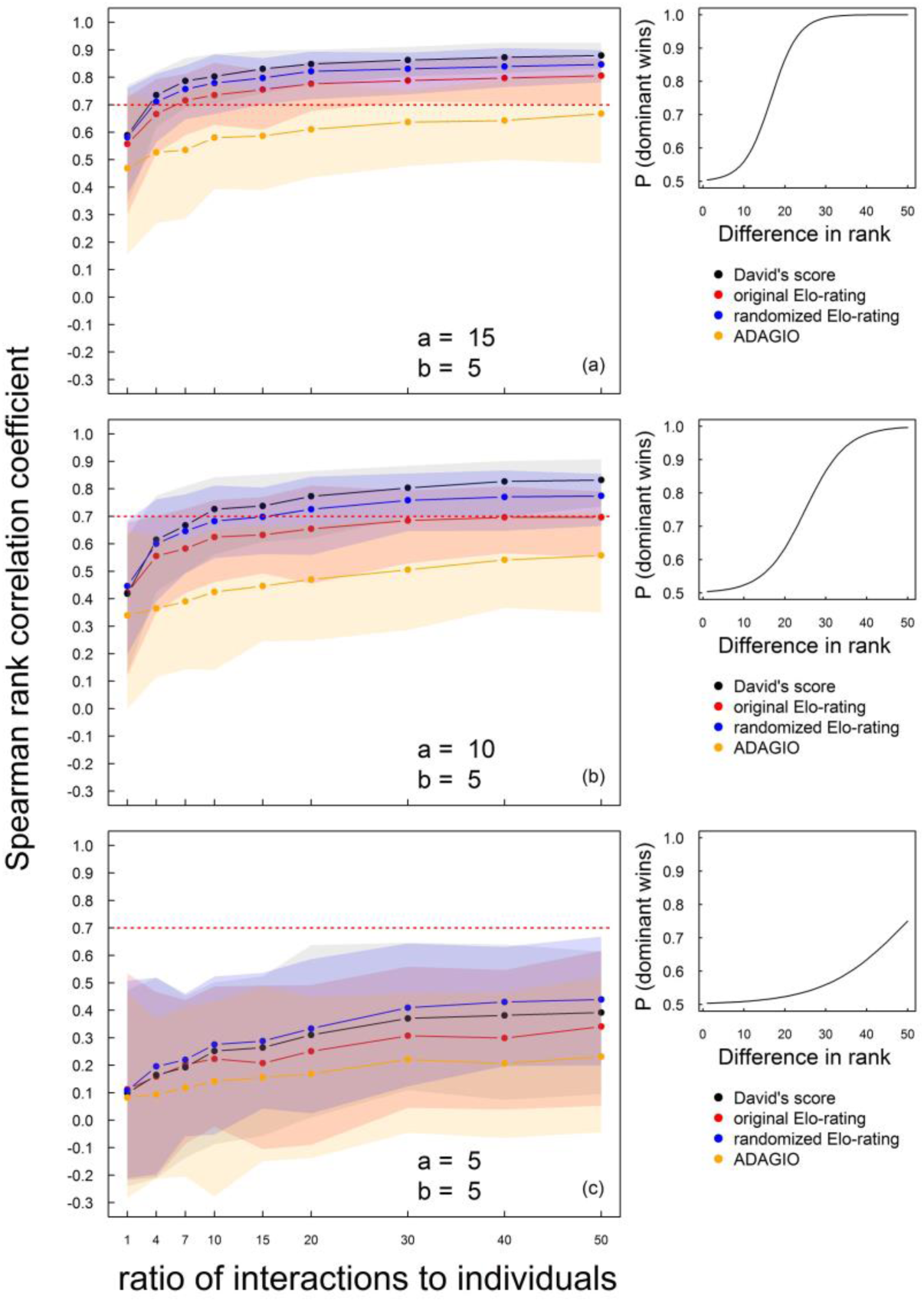
David’s score and the randomized Elo-rating are the two best methods to infer reliable dominance hierarchies. Solid lines and dots represent the mean Spearman rank correlation coefficient (r_S_) between the original and the inferred hierarchy; shading shows the 2.5% and 97.5% quartiles. The red dashed line shows the suggested r_S_ threshold above which inferred hierarchies are highly reliable. The reduced right-hand side panels show the specific hierarchy simulated for generating the interaction datasets. Overall, panels are sorted from intermediate (panels a and b) to very flat hierarchies (panel c). The different hierarchy scenarios shown were created following equation 1.

The results of these simulations can also provide guidance on the number of interactions one should collect to reliably infer dominance hierarchies. Using an r_S_ threshold of 0.70, we suggest to record a ratio of interactions to individuals of 10 to 20 when the real steepness of the hierarchy is unknown – and either David’s score or the randomized Elo-rating should be chosen.

### Hierarchy uncertainty

Last, we explored two easy procedures to estimate hierarchy uncertainty. We first estimated Elo-rating repeatability (Nakagawa & Schielzeth, 2010) using 1 000 individual Elo-ratings obtained from the randomized Elo-rating procedure. We found that, for steep hierarchies, randomized Elo-rating repeatability was high (>0.90 in all cases) and remained relatively stable with the ratio of interactions to individuals (Supplementary Material 3, Fig. S2a-c). For intermediate levels of steepness, randomized Elo-rating repeatability ranged between 0.70 and 0.95 and also remained relatively stable independent of the ratio of interactions to individuals (Fig. 5a,b). In contrast, for very flat hierarchies, randomized Elo-rating repeatability was low and decreased with the ratio of interactions to individuals (Fig. 5c). We therefore suggest a repeatability threshold of 0.70 to differentiate from non-existent/very flat to intermediate/steep hierarchies when enough sampling effort has been done (see method 2 below). Further, our simulations also suggest that the interpretation of the repeatability could include a subsampling routine to determine if repeatability is stable as more data are added (i.e. intermediate/steep hierarchy, Fig. 5a,b) or decreasing (i.e. non-existent/very flat hierarchy, Fig. 5c). That is, the repeatability values provide insights into the steepness of the hierarchy (where higher repeatability scores equate a steeper hierarchy).

**Figure 5.**
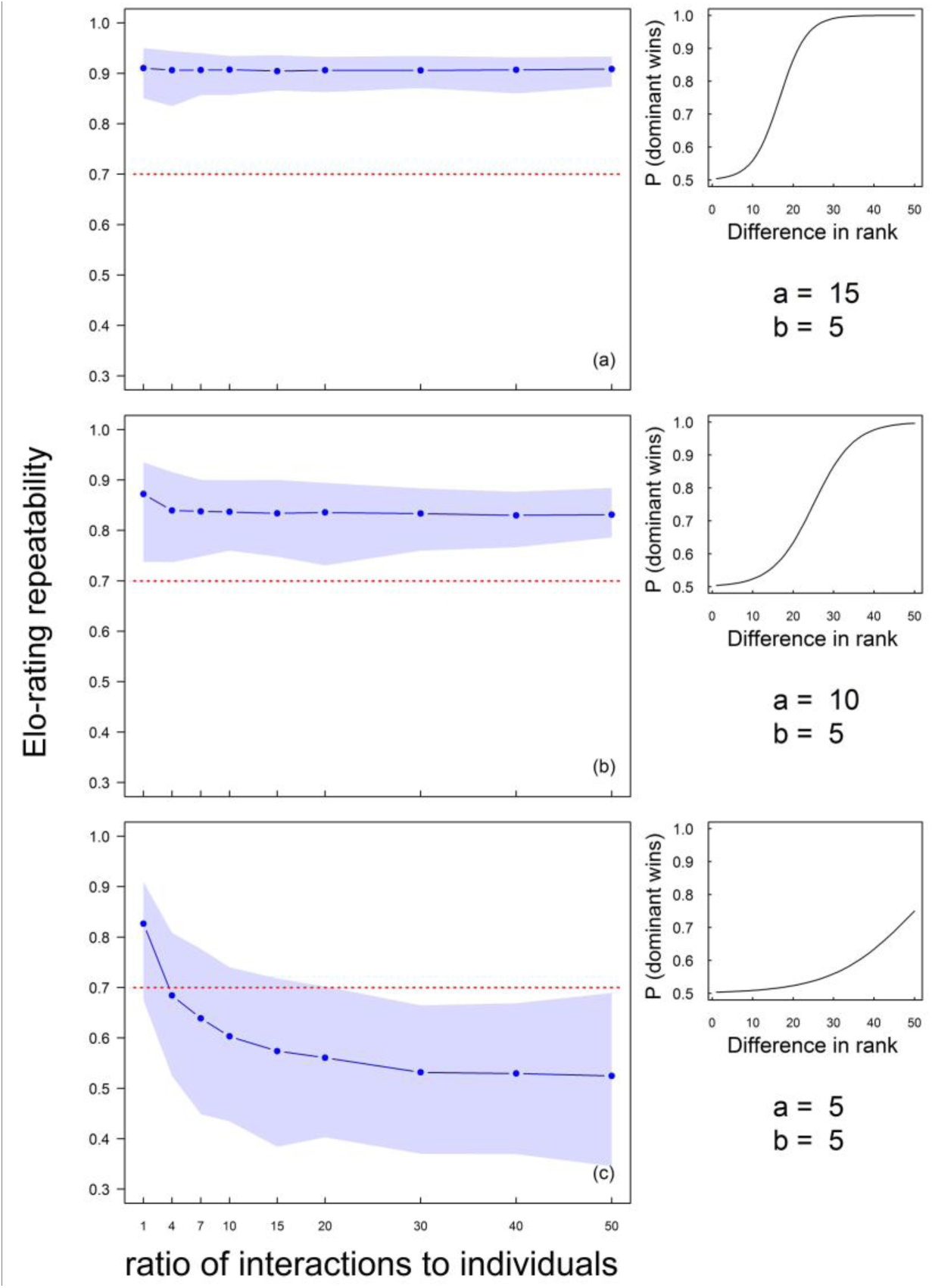
Randomized Elo-rating repeatability increases with the steepness of the hierarchy. Solid lines and dots represent the mean repeatability of the 1 000 individual randomized Elo-ratings estimated for each interaction dataset using the randomized Elo-rating method; shading shows the 2.5% and 97.5% quartiles. The red dashed line shows the suggested repeatability threshold above which inferred dominance hierarchies are highly reliable. The reduced right-hand side panels show the specific hierarchy simulated for generating the interaction datasets. Overall, panels are sorted from intermediate (panel a and b) to very flat hierarchies (panel c). The different hierarchy scenarios shown were created following equation 1.

We propose a second procedure to estimate uncertainty that provides useful information about sampling effort. This method consists of splitting the interaction dataset into two halves and estimating the agreement between the two halves. The Spearman’s rank correlation (r_S_) between the two halves followed a similar logarithmic pattern to the randomized Elo-rating performance, thus also allowing us to make predictions on the level of uncertainty of our measurements and the steepness of the latent hierarchy (Fig. 6). Our results show that, for very steep hierarchies, the r_S_ quickly increased with the ratio of interactions to individuals and stabilized around 0.80 (Supplementary Material 3, Fig. S3a-c). For intermediate hierarchies, however, the r_S_ also increased but did not reach values larger than 0.80 (Fig. 5a,b). For very flat hierarchies, the r_S_ stayed relatively stable and below a threshold value of 0.35 (Fig. 6c). For observational data, adding more interactions into the analysis, and comparing two halves of the interaction dataset, is therefore an informative method for determining whether more sampling effort is required.

**Figure 6.**
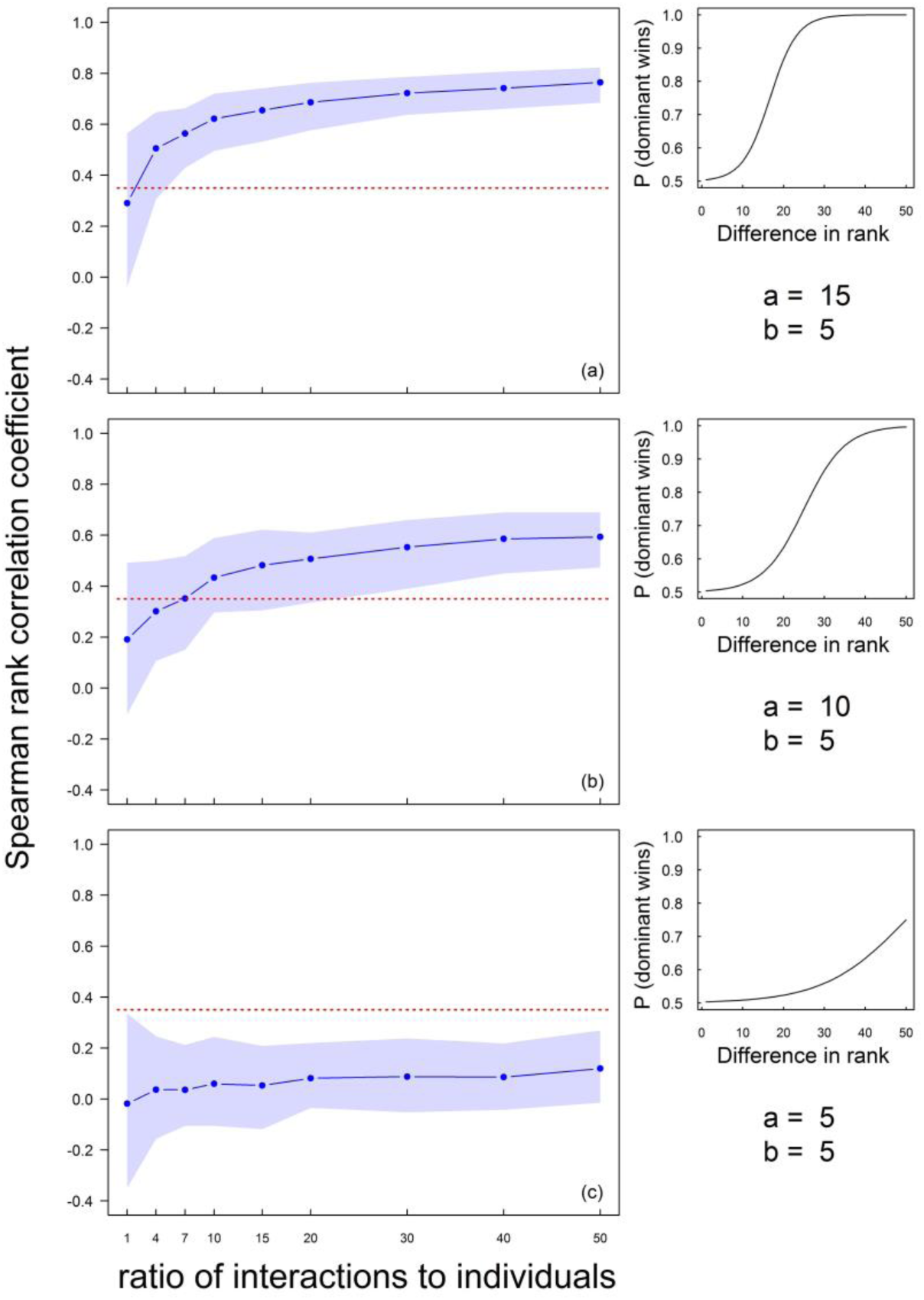
The agreement between the two halves of the interaction dataset increases with the ratio of interactions to individuals and the steepness of the hierarchy. Solid lines and dots represent the mean Spearman rank correlation coefficient (r_S_) between the two halves of the datasets; shading shows the 2.5% and 97.5% quartiles. The red dashed line shows the suggested r_S_ threshold (r_S_ = 0.35) above which inferred dominance hierarchies are highly reliable. The reduced right-hand side panels show the specific hierarchy simulated to generate the interaction datasets. Overall, panels are sorted from intermediate (panel a and b) to very flat hierarchies (panel c). The different hierarchy scenarios shown were created following equation 1.

We conclude that the higher both the randomized Elo-rating repeatability and the r_S_ between the two halves, the steeper the inferred dominance hierarchy is. We further suggest that a repeatability threshold of 0.70 and an r_S_ of 0.35 can be used to differentiate between non-existent/very flat (highly uncertain) and intermediate/steep (certain) hierarchies. Finally, by subsampling the observed data (i.e. recalculating r_S_ with increasingly more data as we have done in Fig. 6) and observing the shape of the resulting metrics is a useful way of determining if the population has been adequately sampled (i.e. adding more data results in very small changes in the r_S_ value).

## DISCUSSION

We tested the performance of four methods to infer dominance hierarchies from dyadic interactions: David’s score (David, 1987), original Elo-rating (Elo, 1978), randomized Elo-rating (this study) and ADAGIO (Douglas et al., 2017). We found that the performance of all methods increases with the steepness of the hierarchy and the ratio of interactions to individuals recorded. We showed that David’s score and the randomized Elo-rating are the two best methods, particularly when hierarchies are not extremely steep. Further, we described two methods for estimating uncertainty in a dataset of observed interactions, and found that uncertainty changes predictably with hierarchy steepness and sampling effort. Based on these results, we provide behavioural and evolutionary ecologists with useful guidelines on the sampling effort necessary to achieve meaningful rank estimations, on how to estimate the uncertainty of their results, and on how to gain insights into the steepness of their hierarchy.

Many methods exist to infer dominance hierarchies from dyadic interaction datasets, and several studies have aimed to assist researchers in choosing the appropriate method (see Introduction). Contrary to most studies, we used simulated interaction datasets rather than real datasets. One of the shortcomings of using real datasets is that the real, latent hierarchy (if existent) is *a priori* unknown and thus, the performance of a method can only be studied relative to other methods, but its reliability cannot be assessed. For example, if all methods suffered the same inherent bias, they could reach similar results but still not reliably infer the real hierarchy. An additional advantage of simulating data is the possibility of generating an incredibly large amount of interaction datasets for each scenario studied, and thus to identify common patterns and gain more general insights on each method. By simulating interaction datasets, we showed that the four methods similarly inferred reliable hierarchies for scenarios of steep hierarchies. In contrast, the methods differed in performance when hierarchies were intermediate or very flat. We showed that David’s score was the best method for intermediate hierarchies, closely followed by the randomized Elo-rating, which also outcompeted its predecessor, the original Elo-rating. Furthermore, as expected, inferred hierarchies were less reliable when using a Poisson process that generates relatively sparse datasets than when using a uniform distribution (Supplementary Material 1, section 1). Contrary to Neumann et al. (2011), our results showed that David’s score still performs better than the original Elo-rating, even when data is relatively sparse. Furthermore, we found that ADAGIO’s performance was sub-optimal. ADAGIO was recently shown to perform better than David’s score and the original Elo-rating (Douglas et al., 2017). The main difference between the study of Douglas et al. (2017) and ours is that the former used Euclidean distances to measure the degree of agreement between original and inferred hierarchies, whereas we used r_S_. While Euclidian distances (i.e. the difference between the real and estimated score) could be a reasonable measure for estimating accuracy in nonlinear hierarchies (Douglas et al., 2017), we tend to prefer relying on rank correlations. One reason for this is that a large population where all individuals have no dominance ranks (e.g. they are all 0) would yield very low Euclidian distances that would suggest very accurate scores despite providing little information in terms of useful dominance data.

Not surprisingly, the performance of all methods increases with hierarchy steepness and the number of interactions recorded. Very steep hierarchies were reliably inferred with a ratio of interactions to individuals of only four (Supplementary Material 3, S1a-c). In contrast, intermediate hierarchies needed a greater sampling effort for inferring reliable dominance hierarchies (Fig. 4). Previous studies already indicated that the social structure of the group studied could affect the method of choice (e.g. Balasubramaniam et al., 2013; Bayly et al., 2006). Here we show the considerable effect that the steepness of the hierarchy has on the performance of the methods studied. Furthermore, many studies have commented on the importance of recording sufficient interactions to infer reliable dominance hierarchies (e.g. Gammell et al., 2003; Neumann et al., 2011). Surprisingly, no clear guidelines existed regarding the sampling effort necessary to infer reliable hierarchies from animals, which in turn has led researchers to apply either untested thresholds to define “sufficient data” (Cole & Quinn, 2011; Devost et al., 2016; Dingemanse & de Goede, 2004; Rat et al., 2015) or no threshold at all (e.g. Hauver, Hirsch, Prange, Dubach, & Gehrt, 2013; Kaburu & Newton-Fisher, 2015). Some studies even failed to report the number of interactions recorded (e.g. Campos & Fedigan, 2013; Flies, Mans, Flies, Grant, & Holekamp, 2016; Stewart & Greives, 2016), impeding researchers to assess the reliability of their results. We suggest that, unless hierarchy is *a priori* known to be very steep, researchers should aim to record a minimum ratio of interactions to individuals of 10 (or ideally 20) to ensure that the dominance hierarchy is reliably inferred. A similar number of interactions was suggested for rating chess players (Glickman & Doan, 2016) but it is considerably larger than other suggestions for animal behaviour (Albers & de Vries, 2001). We acknowledge that recording a ratio of interactions to individuals of 10 to 20 might be challenging for species with low interaction rates such as the red deer (*Cervus elaphus;* Clutton-Brock et al., 1979), but researchers need to be aware of the potential problems of not achieving this threshold, increase (and report) sampling effort whenever possible, and ideally estimate the uncertainty of their dominance data.

Minimizing and estimating measurement error is highly recommended for the study of animal behaviour (Bradshaw, Sims, & Hays, 2007; Martin & Bateson, 2007). Indeed, there is increasing awareness of the need to estimate uncertainty of social data (e.g. Farine & Strandburg-Peshkin, 2015; Lusseau, Whitehead, & Gero, 2008), also, in the study of dominance hierarchies (Adams, 2005). Yet, uncertainty is seldom measured when analysing dominance hierarchies (but see, e.g., Kelstrup, Hartfelder, & Wossler, 2015; Sheppard et al., 2013), possibly because there are no easy-to-use tools to quantify it. Here, we provide two user friendly methods to estimate the level of uncertainty of the inferred hierarchy, and an R package to perform these. First, the randomized Elo-rating method allows calculating individual repeatability as a measure of uncertainty. We have shown that repeatability estimated by randomizing Elo-rating is a good indicator of the steepness of the latent hierarchy, and therefore of the uncertainty of the inferred hierarchy. Furthermore, it is relatively independent of sampling effort, so that, the higher the individual randomized Elo-rating repeatability, both the steeper the latent hierarchy and the more reliable the inferred hierarchy is. We suggest a repeatability threshold of 0.70 to differentiate from non-existent/very flat (highly uncertain) to intermediate/steep (certain) hierarchies. Second, we proposed that uncertainty can also be measured by dividing the interaction dataset in two halves and calculating the level of agreement between the two. Uncertainty measured this way follows a similar pattern to that of method performance, i.e. it decreases with hierarchy steepness and sampling effort. We suggest an r_S_ threshold of 0.35 to differentiate from non-existent/very flat (highly uncertain) to intermediate/steep hierarchies (certain). We conclude that repeatability values and r_S_ below 0.70 and 0.35, respectively, are likely good indicators for a lack of a latent linear dominance hierarchy in the study system.

Additionally, we provide an improvement to the widely accepted original Elo-rating (Elo, 1978). The original Elo-rating is an sequential method proposed for rating chess players that offers some interesting features for the study of animal dominance hierarchies (Albers & de Vries, 2001; Neumann et al., 2011). We show that randomizing 1 000 times the order in which interactions occurred (randomized Elo-rating), and estimating mean individual ranks increases performance compared to the original Elo-rating, particularly when hierarchies are not extremely steep. An important feature of the original Elo-rating is that it allows visualizing hierarchy dynamics (Neumann et al., 2011), this feature is not possible when using matrix-like methods such as David’s score. Visualizing hierarchy dynamics might be important to study social dynamics (Neumann et al., 2011) and/or winner-loser effects (Hsu & Wolf, 1999). Nonetheless, the randomized Elo-rating is useful when researchers have a good reason to believe that the dynamics are relatively stable and rank an inherent property of the individual. Further, we suggest that researchers can control for factors such as changing ranks or winner-loser effects by controlling which parts of the data are randomised. For example, researchers interested in tracking how individual rank changes across days, could increase the precision of the daily rank estimates by randomizing observations within each day. The randomized Elo-rating further allows easy estimation of uncertainty for individual Elo-ratings or ranks. For example, as discussed above, researches can estimate Elo-rating repeatability, however, they can also obtain common measures of dispersion such as confidence intervals or standard deviation for each individual. So far, this was only possible using the Bayesian procedure proposed by Adams (2005). Finally, we believe that some researchers avoid the Elo-ratings if they have recorded their data in a matrix format (and thus do not have the sequence of the interactions). We suggest that in these cases, the randomized Elo-rating would avoid the potential spurious results that could arise by assigning a single random order to the observations.

Finally, we acknowledge and discuss some of the potential limitations of this study. First, due to computational limitations, we only investigated four methods. We however provided an R package and the R code used in this study to aid researchers interested in exploring other methods (see Sánchez-Tójar et al., 2017). Second, one of the limitations of the randomized Elo-rating is that it does not include tied or undecided interactions. Undecided interactions are very rare (1.8% of all interactions from 40 empirical interactions datasets; McDonald & Shizuka, 2013), and researchers not always agree on how to interpret them (e.g. Balasubramaniam et al., 2013). In fact, it could be argued that part of the undecided interactions are just so because of the difficulties of understanding animal behaviour. We therefore suggest that undecided interactions should always be reported but not necessarily used when inferring dominance hierarchies. Third, in this study we focused on inferring dominance hierarchies, i.e. we focus on ranks. However, David’s score, and the original and randomized Elo-ratings were originally developed for estimating individual indices of (fighting) success. We believe that our suggestions can further help researchers aiming to study individual (fighting) success rather than dominance hierarchies.

### Conclusions

We have shown how sampling effort and the steepness of the underlying hierarchy (a latent feature *a priori* unknown by the researcher) affect method performance. We have suggested and provided with a new method, the randomized Elo-rating (R package “aniDom”: Farine & Sánchez-Tójar, 2017). We have shown that David’s score and the randomized Elo-rating are the two best methods to infer linear dominance hierarchies, particularly when the latent hierarchy is not extremely steep. Furthermore, we have introduced two easy procedures to evaluate hierarchy uncertainty at both the individual and the group level. Last but not least, we have provided clear guidelines on how much sampling effort is required to infer reliable hierarchies. We believe that the procedures outlined here are simple to implement and that the guidelines we provide will help researchers aiming to study dominance hierarchies. Finally, we hope that this work will help mitigating some of the problems recently raised (Nakagawa & Parker, 2015) in the broader field of behavioural research.

## ACKNOWLEDGEMENTS

AST is member of the International Max Planck Research School (IMPRS) for Organismal Biology. We thank Antje Girndt for constructive feedback on the manuscript. AST and JS were funded by the Volkswagen Foundation. The authors declare that they have no conflict of interest.

## RESOURCES

We provide all of the source code used for our data (Sánchez-Tójar, Schroeder, & Farine, 2017). We also provide with a free R package to run our implementations (“aniDom”: Farine & Sánchez-Tójar, 2017). Further, we note that our implementation of the original non-randomized Elo-rating outperforms existing R packages for some scenarios (Sánchez-Tójar, Schroeder, & Farine, 2017).

